# Effect of methylene blue on the formation of oxidized phospholipid vesicles

**DOI:** 10.1101/403634

**Authors:** J-F. Fabre, M. Cerny, A. Cassen, Z. Mouloungui

## Abstract

Soybean phosphatidylcholine, which is rich in linoleic acid, was oxidized with singlet oxygen through photosensitization with methylene blue. This compound facilitates the oxidation of phospholipids relative to the reaction with free unsaturated fatty acids. A response surface methodology was used to control oxidation, with methylene blue concentration and the amount of available air as independent variables. The conjugated diene-to triene ratio was then monitored. Hydroperoxide yield dependent principally on the amount of air, whereas photosensitizer concentration strongly influenced the size and zeta potential of vesicles formed by the sonication of oxidized phospholipids in water. Methylene blue plays an important role in the surface charge expression and ion permeability of these vesicles.

## 1. Introduction

Phospholipids were initially considered to be a by-product of oil refining. However, over time, these amphipathic molecules have become essential components of food, cosmetic and pharmaceutical formulations. Their organization as liposomes in water confers considerable added value for drug encapsulation and delivery(1–3). The size, shape and permeability of liposomes depend strongly on the nature and composition of the phospholipids they contain. For example, liposomes formed with phosphatidylcholine (PC), the most abundant phospholipid in animals and plants, are generally larger than those formed with phosphatidylethanolamine(4). The nature of the fatty acids is also of great importance. Saturated fatty acids form more compact membranes that are generally less permeable for some compounds than the equivalent unsaturated fatty acids(5–7) but, for some ions, the membranes formed with unsaturated fatty acids are less permeable than those formed with saturated fatty acids(8). Long fatty acid chains generally result in lower permeability to ions(9, 10). Phospholipid oxidation, which occurs at double bonds, can have several effects. Oxidation can stimulate lipid flip-flop within the membrane(11, 12), and oxidized liposomes are generally considered more permeable to water (13, 14), glucose(15) and dextran(16). High levels of oxidation can also lead to pore formation (17, 18, 19), and membrane disruption, mostly due to the presence of shortened carboxyacyl units (20), as hydroperoxides cause less damage. However, all these features are heavily dependent on the nature and concentration of the oxidized compounds and it is thus essential to control oxidation, to modify the hydrophilic and permeation properties of these liposomes as desired. Soybean seeds are one of the most widely used sources of commercial phospholipids. Phosphatidylcholine is the principal soybean phospholipid. Its fatty acid chains are mostly unsaturated (oleic, linoleic, linolenic acids), making it a good model for oxidation studies. A number of different methods are available for oxidizing soybean phospholipids(21, 22) but the use of singlet oxygen is a simple and powerful approach. This method generally involves the disproportionation of hydrogen peroxide to generate singlet oxygen, and the use of molybdate as a catalyst (23, 24).

However, a photosensitizer can also be used, and this method is often used with fatty acids or phospholipids. Various dyes can be used for this purpose, including protoporphyrin IX (25), rose bengal (26), pheophorbide (27) and methylene blue (MB), one of the most widely used photosensitizers (28–32). Under illumination, the methylene blue cation reaches an excited singlet state. As a conjugated dye containing a single sulfur as a ring heteroatom, it can undergo efficient intersystem crossing to reach a triplet state with a relatively long lifetime. Photooxidation can then occur through two possible mechanisms. In type II mechanisms, triplet oxygen reacts with MB through an energy transfer process to form singlet oxygen, while the methylene blue returns to its fundamental level (33, 34). Singlet oxygen, which is highly electrophilic, generates hydroperoxides on reaction with unsaturated chains. In type I mechanisms, MB reacts with the substrate via electron transfer to form radicals. Hydroperoxides are generally formed first, through a type II mechanism, which is then followed by a type I mechanism (35). However, the contribution of methylene blue is often limited to the production of singlet oxygen, and this compound is generally removed from the reaction medium for the purification of oxidized products. To our knowledge, its possible contribution to the formation and properties of vesicles has hardly been explored.

## 2. Materials and methods

Methylene blue trihydrate, linoleic acid (>99%), linolenic acid (>99%), and oleic acid (99%), for use as analytical reactants and standards, were purchased from Sigma Aldrich (St Quentin Fallavier, France). Soybean lecithin (ca. 90% phosphatidylcholine) was purchased from VWR (Fontenay sous Bois, France). Dilinoleylphosphatidylcholine (>99%) was purchased from COGER (Paris, France)

### Chemical analysis

The fatty acid profile of the phospholipids was determined after *trans*-methylation with TMSH (0.2 M trimethylsulfonium hydroxide in methanol) according to AFNOR Method NF EN ISO 12966-3. The fatty acid methyl esters (FAME) obtained by this transesterification reaction were analyzed with a gas chromatograph (36) equipped with a CP-select CB column (50 m long, 0.32 mm i.d., 0.50 µm film thickness); helium was used as the carrier gas, at a flow rate of 1.2 mL/minute; the split injector (1:100) and FID were maintained at 250°C; the initial oven temperature was set to 185°C for 40 minutes, increased to 250°C at a rate of 15°C/minute and maintained at this temperature for 10 minutes.

NMR experiments were carried out on a Bruker AVANCE III HD NMR spectrometer operating at 500.13 MHz for 1 h. This machine was equipped with a 5 mm BBFO ATMA Prodigy cryoprobe. This cryoprobe enhances the probe sensitivity at room temperature by a factor of 2 to 3 for X-nuclei from 15N to 31P. NMR experiments were recorded at 298 K, with the zgig30 pulse sequence. The recycle delay was adjusted to 2 s and the number of scans was set to 16. Topspin 3.2 software was used for data acquisition and processing in all NMR experiments.

The conjugated dienes and trienes generated by phospholipid oxidation were characterized by spectrophotometry with a SHIMADZU UV1800 spectrophotometer, in quartz cuvettes. The reference cuvette contained absolute ethanol. The contribution of methylene blue to the absorbances at 233 nm and 290 nm was ignored.

### Photosensitization

Solutions of different concentrations of methylene blue in absolute ethanol were prepared in amber bottles and stored in the dark. Phospholipids were dissolved in these solutions at a concentration of 20 mg/mL, and the resulting solutions were illuminated for eight hours in 20 mL glass tubes (diameter: 15 mm) with an ATLAS SUNTEST CP+ device equipped with a 1500 W xenon lamp simulating a spectral distribution close to that of natural sunlight.

### Size and zeta potential measurements

The size and zeta potential of phospholipid vesicles were analyzed with a NANOSIZER ZS granulometer (Malvern). A 1 mL sample of the 20 mg/mL phospholipid solution in absolute ethanol was dried under nitrogen. The resulting film was diluted in 2 mL of distilled water and sonicated for 3 minutes with a laboratory ultrasound probe (SONICS VIBRACELL −500 equipped with a 3mm diameter probe, pulsation mode: 5 s ON/10 s OFF, amplitude: 57µm/20%). The temperature was allowed to stabilize at 25°C for one minute, and measurements were performed three times, considering a refractive index for the lipid bilayer of 1.45 (37) and an absorption factor of 0.001. A backscattering detector was positioned at 173°. Zeta potential values were obtained by correlating mobility measurements with the Smoluchowsky model.

### Permeation studies

If the solute concentration differs between the phase within vesicles and the external phase, then there is an osmotic pressure exerts at both sides. Vesicles may undergo water or salt permeation to equilibrate this pressure. If selective, this permeation leads to a change in volume. For example, volume decreases if the vesicle rejects water to increase its internal solute concentration, or increases if the vesicle accepts water to decrease its internal solute concentration. Absorbance measurements are a good tool for measuring such changes, provided that they occur rapidly. Indeed, the absorbance of a stable dispersed medium is highly dependent on the refractive index of the dispersed particles, their concentration and their size. For a given refractive index and vesicle concentration, the change in absorbance is related to changes in size and, thus, to the exchanges of water and solutes between the vesicles and the surrounding medium.

According to Mie theory of light scattering (38), for a given refractive index, solutions of nanometric particles (10-100 nm) scatter much less light than solutions of micrometric particles, and have a lower light extinction value. The initial absorbance of vesicle solutions (before the occurrence of osmotic changes) may, therefore, be directly linked to vesicle size. It is difficult to measure absorbance at the precise time at which the vesicle suspension is dispersed in the medium. The absorbance reached at equilibrium (or at infinite time) is therefore more usually measured. This approach also has the advantage that the delayed absorbance variation generally reflects a more precise equilibration of osmotic pressures. Decreases in absorbance have generally been linked to a global swelling, whereas shrinkage is accompanied by an increase in absorbance (6, 39).

The variation of absorbance can be modeled with an exponential law (see supporting information):

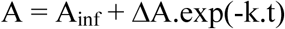

where A_inf_ is the absorbance at infinite time, ΔA is the change in absorbance between the initial measurement and infinite time, and k is a constant.

The swelling and shrinkage of phospholipid structures in hypertonic solutions were, therefore, measured with a Spectrostar NANO spectrophotometer, at a fixed temperature of 25°C, in polypropylene cuvettes. Kinetic measurements of extinction at 400 nm were performed at five-second intervals.

### Response surface methodology

The experiment was designed and analyzed with NEMRODW 2000 (Mathieu D, Nony J, Phan-Tan-Luu R. NEMROD-W software. LPRAI; Marseille: 2000).

The experimental values were regressed with a second-order polynomial model:

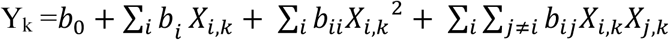

Where *Y*_*k*_ is the calculated response value in the *k*^th^ experiment, *X*_*i,k*_ is the coded variable *i* for the *k*^th^ experiment, *b*_*0*_ is the intercept term, *b*_*i*_ is the main coefficient for each variable, *b*_*ii*_ is the squared coefficient and *b*_*ij*_ is the interaction term. The pertinence of the model was assessed with the Fisher-Snedecor test, comparing model and residual variances, and its validity (or descriptive importance) was determined by comparing the residual and experimental variances (see supplementary information). A proba.1 value below 5 was considered to indicate pertinence of the model (at a 95% confidence level). A proba.2 value below 5 was interpreted as indicating that the model did not describe the observed values (little chance that any deviation from the model could be explained by the experimental error). Measurements were also performed for test points, to assess the predictive value of the model in the experimental domain.

## 3. Results and Discussion

### 3.1. Fatty acid distribution and oxidation

The fatty acid distribution of soybean phosphatidylcholine can be obtained both by ^1^H NMR analysis (40, 41) on intact phospholipids and by GC after transmethylation. NMR analysis is based on the different chemical shifts between the terminal methyl, allyl and alkene hydrogens of oleic, linoleic and linolenic acids. It is more difficult to determine the composition of saturated acids by this method, which is therefore used mostly to determine unsaturated fatty acid content. The two methods gave similar results for unsaturated fatty acid composition (table 1).

**Table 1.** Fatty acid distribution of soybean phosphatidylcholine according to GC and ^1^H NMR analyses

| Fatty acid | % by GC | % by NMR |
| --- | --- | --- |
| C16:0 – palmitate | $14.11 \pm 0.15$ | <19 |
| C18:0 – stearate | $3.85 \pm 0.13$ | <5 |
| C18:1 – oleate | $11.31 \pm 0.03$ | 13.7 |
| C18:1n-7c – vaccenate | $1.66 \pm 0.06$ | |
| C18:2n-6c – linoleate | $63.27 \pm 0.14$ | 59.5 |
| C18:3 – linolenate | $5.80 \pm 0.05$ | 7.5 |

NMR cannot distinguish between oleate and vaccenate. The results obtained indicate that the principal fatty acids of soybean PC were linoleic, palmitic, then oleic and linolenic acids. The fatty acids of this molecule are, therefore, mostly polyunsaturated.

The oxidation of soybean PC was further investigated by preparing solutions of free linoleic, linolenic, and oleic acids and soybean PC, at a concentration of 20 mg/mL, in absolute ethanol. A 10 mL sample of each solution was poured into a 20 mL glass tube, which was then hermetically sealed. The tube was then illuminated for 8 hours, and NMR analysis was performed on the products. The oxidation of linoleic, linolenic and oleic acids was characterized by the appearance of peaks (fig. 1) between 4 and 4.5 ppm (C<u>H</u>-OOH), 5.4 and 6.5 ppm (conjugated double bonds) and 11 and 11.5 ppm (-OO<u>H</u>)(42, 43, 44), as shown in the supplementary information.

**Figure 1.**
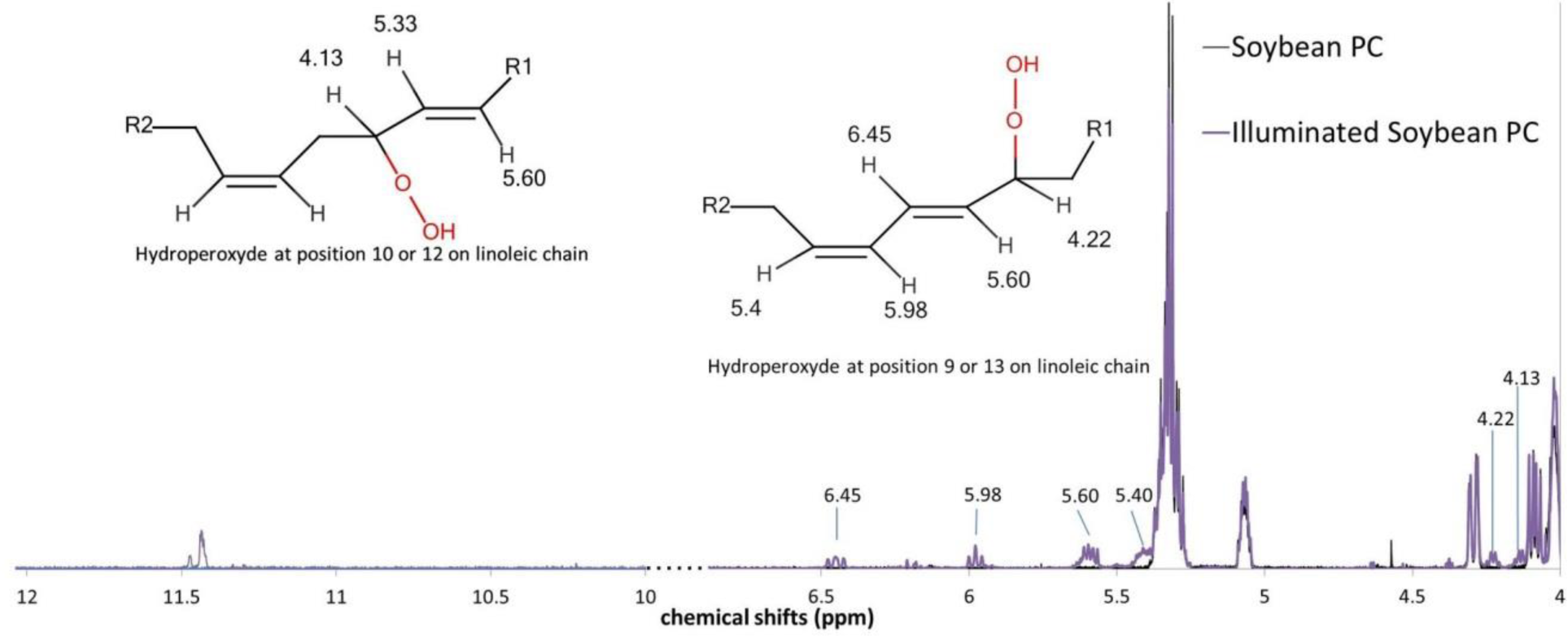
Difference between the ^1^H-NMR spectra of soybean phosphatidylcholine and oxidized phosphatidylcholine (the spectra have been truncated to highlight the main differences between them)

The various hydroperoxide signals can be integrated to determine the formation yield. As for fatty acids (table 2), similar integration results were obtained if we considered both hydrogens of the hydroperoxide group (-C<u>H</u>OO<u>H</u>). The protons carried by the carbon in soybean phospholipids are partially screened by peaks corresponding to the polar phosphocholine moiety. Only the hydrogens corresponding to linoleic acid are visible. Integration is therefore best performed with the hydrogen carried by the oxygen (–CHOO<u>H).</u>

**Table 2.**
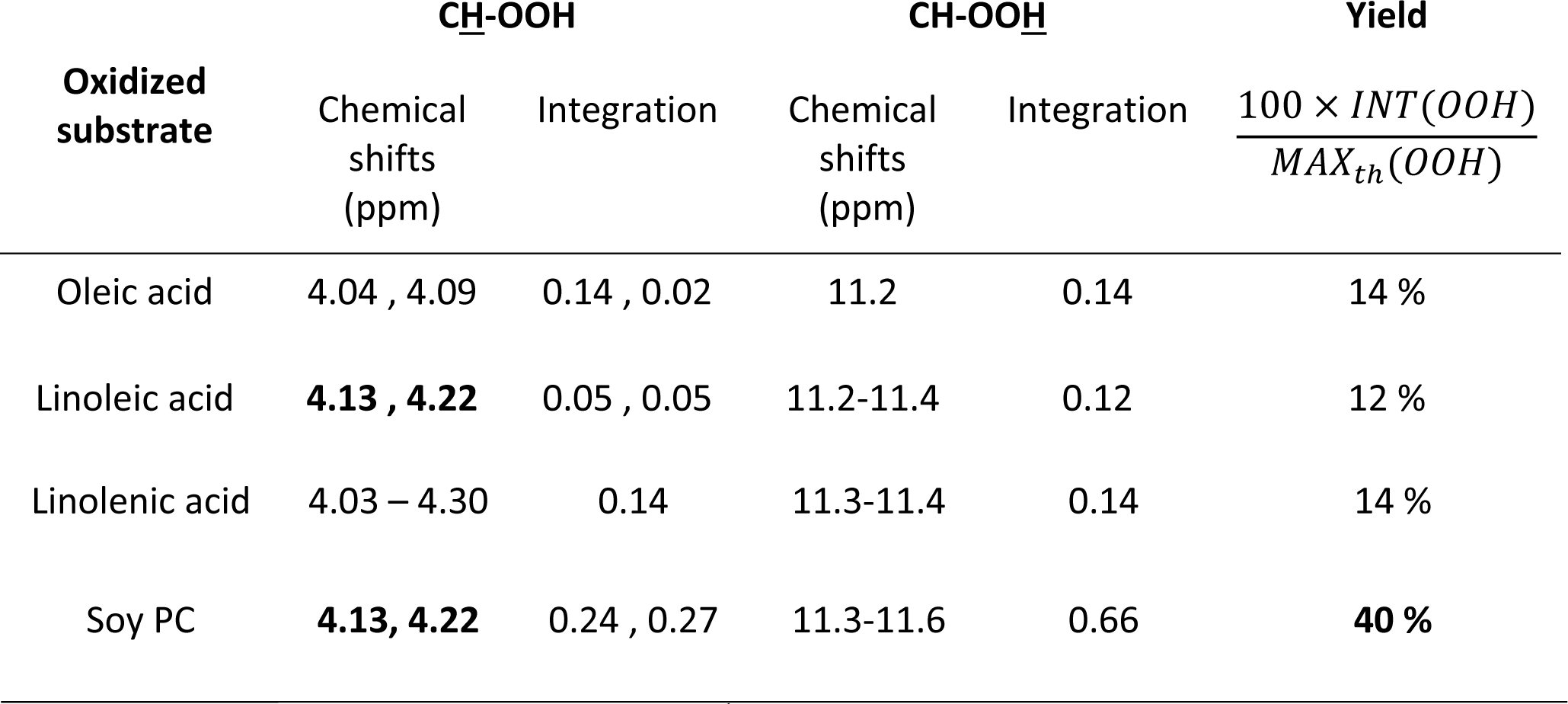
Hydroperoxide yield according to ^1^H NMR analysis

Knowing the composition of the phospholipid tested, which consisted of 80% unsaturated fatty acids, we can calculate that, if one hydroperoxide per unsaturated chain is considered, the maximum amount of hydroperoxide in one molecule of phospholipid would be 1.6. For soybean phospholipid, the hydroperoxide yield can, therefore, be expressed as: 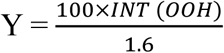 where INT(OOH) is the integration value for OO<u>H</u>/molecule

Yields were similar and low for the three unsaturated fatty acids (table 2), whereas soybean phospholipid was much more oxidized. The oxidation of linoleic and linolenic acids can also be monitored by spectrophotometry. Hydroperoxide generation leads to the formation of conjugated double bonds the absorbance of which can be monitored over time. Conjugated dienes absorb at 230-235 nm, whereas conjugated trienes absorb at 270-275 nm. The absorbances of the phospholipid solutions were measured at different dilutions, to obtain the extinction coefficient with the Beer-Lambert formula:

A = ε_n_.c_n_.l or A= ε_m_.c_m_.l regardless of whether molar concentration or mass concentration is considered.

*ε*_*n*_ is the molar extinction coefficient (L.mol^−1^.cm^−1^), *c*_*n*_ is the molar concentration (mol.L^−1^), *l* is the path length (cm) in the solution, *ε*_*m*_ is the mass extinction coefficient (L.g^−1^.cm^−1^), and *c*_*m*_ is the mass concentration (g.L^−1^). For the comparison of molar extinction coefficients (table 3), an approximate molecular weight of 782 g/mol was chosen for soybean phosphatidylcholine, corresponding to the molecular weight of dilinoleylphosphatidylcholine.

**Table 3.** Extinction coefficients of soybean PC and its polyunsaturated fatty acid constituents

| Oxidized substrate | $\varepsilon_c$ | $\varepsilon_c$ | Approx. $\varepsilon_n$ | Approx. $\varepsilon_n$ |
| --- | --- | --- | --- | --- |
|  | (233nm) | (270nm) | (233nm) | (270nm) |
| | ($\text{L} \cdot \text{g}^{-1} \cdot \text{cm}^{-1}$) | ($\text{L} \cdot \text{g}^{-1} \cdot \text{cm}^{-1}$) | ($\text{L} \cdot \text{mol}^{-1} \cdot \text{cm}^{-1}$) | ($\text{L} \cdot \text{mol}^{-1} \cdot \text{cm}^{-1}$) |
| Soy PC | 5.88 | 0.19 | 4200 | 140 |
| Linoleic acid | 5.38 | No peak | 1450 | No peak |
| Linolenic acid | 7.94 | 0.9 | 1650 | 235 |

Despite their similar hydroperoxide yields, oxidized linolenic acid had a higher extinction coefficient than linoleic acid at 233 nm, which can be explained by the larger number of unsaturated bonds, resulting in a larger number of positions at which conjugated dienes can be produced in linolenic (positions 9, 12, 13, 16) than in linoleic (positions 9 and 13) acids. Soybean phospholipid had a higher mass extinction coefficient than linoleic acid, and a molar extinction coefficient more than twice those of linoleic and linolenic acids. Its extinction coefficient at 270 nm was also less than half that of linolenic acid, with linolenic acid accounting for less than 10% of its composition. Soybean phospholipid is, thus, much more easily oxidized than its constitutive unsaturated fatty acids, consistent with the NMR results. Methylene blue may, therefore, have a higher affinity for phospholipids than for fatty acids.

### 3.2. Response surface methodology

Singlet oxygen production depends principally on the amount of photosensitizer and available triplet oxygen, provided that the samples are sufficiently illuminated. It was decided to work with air rather than pure oxygen, to obtain a procedure that was both simplified and scalable. For a constant temperature of 25°C, a change in the relative humidity of the air of less than 10% during sample tube filling and a constant atmospheric pressure, the oxygen content of air can be considered stable and the amount of oxygen in the sample tubes is linearly correlated with the volume of available air. Using the highest illumination power available, >750 W/m^2^ in the range 300-800nm, and limiting the duration of illumination to eight hours, the photosensitizer concentration and the volume of the sample solution in the sealed tube were varied, to modify the amount of air available. Phospholipid concentration was maintained at 20 mg/mL. The aim was determine whether several valuable responses could be described qualitatively and/or quantitatively within a large domain of variations. Five values of log (1/V) were chosen for the volume (V) of the phospholipid solution, and three values of log(C) for the concentration of methylene blue.

A Doehlert matrix was used, with seven minimal experiments, two repetitions at the center of the domain, one additional repetition and four test points at different coordinates (table 4).

**Table 4.** Extended Doehlert matrix for controlled singlet oxygen oxidation

| EXP. | V<br>(mL) | [MB]<br>(mol.L<sup>-1</sup>) | LOG(1/V)<br>real value | LOG ([MB])<br>real value | LOG(1/V)<br>coded value<br>X1 | LOG ([MB])<br>coded value<br>X2 |
| --- | --- | --- | --- | --- | --- | --- |
| 1 | 2 | 1x10<sup>-4</sup> | -0.3 | -4 | 1 | 0 |
| 2 | 20 | 1x10<sup>-4</sup> | -1.3 | -4 | -1 | 0 |
| 3 | 3.5 | 1x10<sup>-3</sup> | -0.55 | -3 | 0.5 | 0.866 |
| 4 | 11 | 1x10<sup>-5</sup> | -1.05 | -5 | -0.5 | -0.866 |
| 5 | 3.5 | 1x10<sup>-5</sup> | -0.55 | -5 | 0.5 | -0.866 |
| 6 | 11 | 1x10<sup>-3</sup> | -1.05 | -3 | -0.5 | 0.866 |
| 7 | 6 | 1x10<sup>-4</sup> | -0.8 | -4 | 0 | 0 |
| 8 repetition | 6 | 1x10<sup>-4</sup> | -0.8 | -4 | 0 | 0 |
| 9 repetition | 6 | 1x10<sup>-4</sup> | -0.8 | -4 | 0 | 0 |
| 10 repetition | 11 | 1x10<sup>-3</sup> | -1.05 | -3 | -0.5 | 0.866 |
| 11 test point | 6 | 1x10<sup>-3</sup> | -0.8 | -3 | 0 | 0.866 |
| 12 test point | 6 | 1x10<sup>-5</sup> | -0.8 | -4 | 0 | -0.866 |
| 13 test point | 15 | 1x10<sup>-3</sup> | -1.18 | -3 | -0.76 | 0.866 |
| 14 test point | 15 | 1x10<sup>-5</sup> | -1.18 | -5 | -0.76 | -0.866 |

An additional 6 mL sample of solution with a methylene blue concentration of 1×10^−4^ M was covered with a sheet of aluminum foil to block out the light as a control, and another control consisted of methylene blue in ethanol alone. The color of these two control samples was unchanged after 8 h in simulated sunlight, indicating that the methylene blue was not degraded in these conditions. By contrast, the blue color of all the other samples in the experimental design decreased, indicating a degradation of the methylene blue in the presence of light, oxygen and phospholipids.

#### 3.2.1 Yield and extinction coefficients

The hydroperoxide yield for all experiments was determined by integrating the NMR signals corresponding to hydroperoxide (-OOH). The analysis of variance yielded a Proba.1 value <0.01, indicating that the regression was highly significant (confidence level of 99.9%). However, a comparison of repeat experiments (table 5) revealed a small experimental error that was insufficient to account for the difference between the experimental and calculated values (Proba.2 =0.041). The model can provide qualitative information. An examination of the different values of the experimental points (fig. 2a) and the correlation table (table 6) clearly showed that yield was strongly dependent on the first factor, log(1/V).

**Table 5.** Observed and predicted values for the various dependent variables measured.

| Exp. | X1 | X2 | Yield<br>obs.<br>% | Yield<br>pred.<br>% | $\epsilon_{233}$<br>obs.<br>L.g<sup>-1</sup>.cm<sup>-1</sup> | $\epsilon_{233}$<br>pred.<br>L.g<sup>-1</sup>.cm<sup>-1</sup> | $\epsilon_{233}/\epsilon_{270}$<br>obs. | $\epsilon_{233}/\epsilon_{270}$<br>pred. | Z<br>obs.<br>nm | Z<br>pred.<br>nm | $\zeta$<br>obs.<br>mV | $\zeta$<br>pred.<br>mV | A<sub>inf</sub><br>obs. | A<sub>inf</sub><br>pred. |
| --- | --- | --- | --- | --- | --- | --- | --- | --- | --- | --- | --- | --- | --- | --- |
| 1 | 1 | 0 | 63.13 | 69.21 | 13.89 | 15.00 | 27.54 | 27.86 | 146 | 141 | -0.03 | -6.13 | 0.114 | 0.113 |
| 2 | -1 | 0 | 26.25 | 20.17 | 5.20 | 4.10 | 15.96 | 15.64 | 338 | 342 | 4.08 | 10.18 | 0.409 | 0.410 |
| 3 | 0.5 | 0.866 | 60.63 | 54.55 | 15.91 | 14.81 | 16.17 | 15.85 | 126 | 130 | 5.24 | 11.34 | 0.138 | 0.193 |
| 4 | -0.5 | -0.866 | 30.63 | 36.71 | 4.54 | 5.65 | 28.59 | 28.91 | 523 | 518 | 2.54 | -3.56 | 0.253 | 0.252 |
| 5 | 0.5 | -0.866 | 43.13 | 43.96 | 6.74 | 5.64 | 27.71 | 27.39 | 308 | 312 | 2.55 | 8.65 | 0.190 | 0.191 |
| 6 | -0.5 | 0.866 | 2.50 | 5.86 | 3.24 | 3.89 | 1.98 | 2.11 | 134 | 126 | 47.60 | 39.85 | 0.362 | 0.376 |
| 7 | 0 | 0 | 43.75 | 43.96 | 9.02 | 8.81 | 29.77 | 32.31 | 228 | 237 | 4.31 | 3.54 | 0.163 | 0.161 |
| 8 | 0 | 0 | 45.00 | 43.96 | 8.70 | 8.81 | 32.80 | 32.31 | 248 | 237 | 3.33 | 3.54 | 0.166 | 0.161 |
| 9 | 0 | 0 | 43.13 | 43.96 | 8.72 | 8.81 | 34.36 | 32.31 | 236 | 237 | 2.97 | 3.54 | 0.155 | 0.161 |
| 10 | -0.5 | 0.866 | 3.13 | 5.86 | 3.44 | 3.89 | 1.91 | 2.11 | 122 | 126 | 38.20 | 39.85 | 0.391 | 0.376 |
| 11 | 0 | 0.866 | 26.25 | 30.02 | 8.23 | 9.17 | 4.25 | 11.62 | 96 | 127 | 26.00 | 25.97 | 0.169 | 0.233 |
| 12 | 0 | -0.866 | 40.00 | 36.70 | 6.21 | 5.46 | 29.71 | 30.79 | 284 | 414 | -1.72 | 2.92 | 0.231 | 0.197 |
| 13 | -0.76 | 0.866 | 0.63 | -6.57 | 2.64 | 1.30 | 1.52 | -4.93 | 122 | 126 | 42.50 | 46.77 | 0.546 | 0.470 |
| 14 | -0.76 | -0.866 | 16.25 | 36.86 | 2.72 | 5.89 | 20.06 | 25.85 | 360 | 573 | -2.30 | -7.23 | 0.414 | 0.300 |

**Table 6.** Coefficients of correlation between different variables.

|  | *X1* | *X2* |  |  |  |  |  |  |  |
| --- | --- | --- | --- | --- | --- | --- | --- | --- | --- |
| | $\log(1/V)$ | $\log(C)$ | Yield | $\epsilon_{233nm}$ | $\epsilon_{233nm}/\epsilon_{270nm}$ | Z average | $\zeta$ | $A_{inf}$ | |
| *X1* | 1 |  |  |  |  |  |  |  |  |
| *X2* | -0.06 | 1 |  |  |  |  |  |  |  |
| Yield | 0.82 | -0.31 | 1 |  |  |  |  |  |  |
| $\epsilon_{233nm}$ | 0.83 | 0.14 | 0.89 | 1 | | | | | |
| $\epsilon_{233nm}/\epsilon_{270nm}$ | 0.44 | -0.74 | 0.75 | 0.39 | 1 | | | | |
| Z average | -0.29 | -0.85 | 0.09 | -0.31 | 0.56 | 1 |  |  |  |
| $\zeta$ | -0.38 | 0.76 | -0.77 | -0.45 | -0.88 | -0.60 | 1 | | |
| $A_{inf}$ | -0.84 | 0.17 | -0.87 | -0.81 | -0.63 | 0.03 | 0.56 | 1 | |

**Figure 2.**
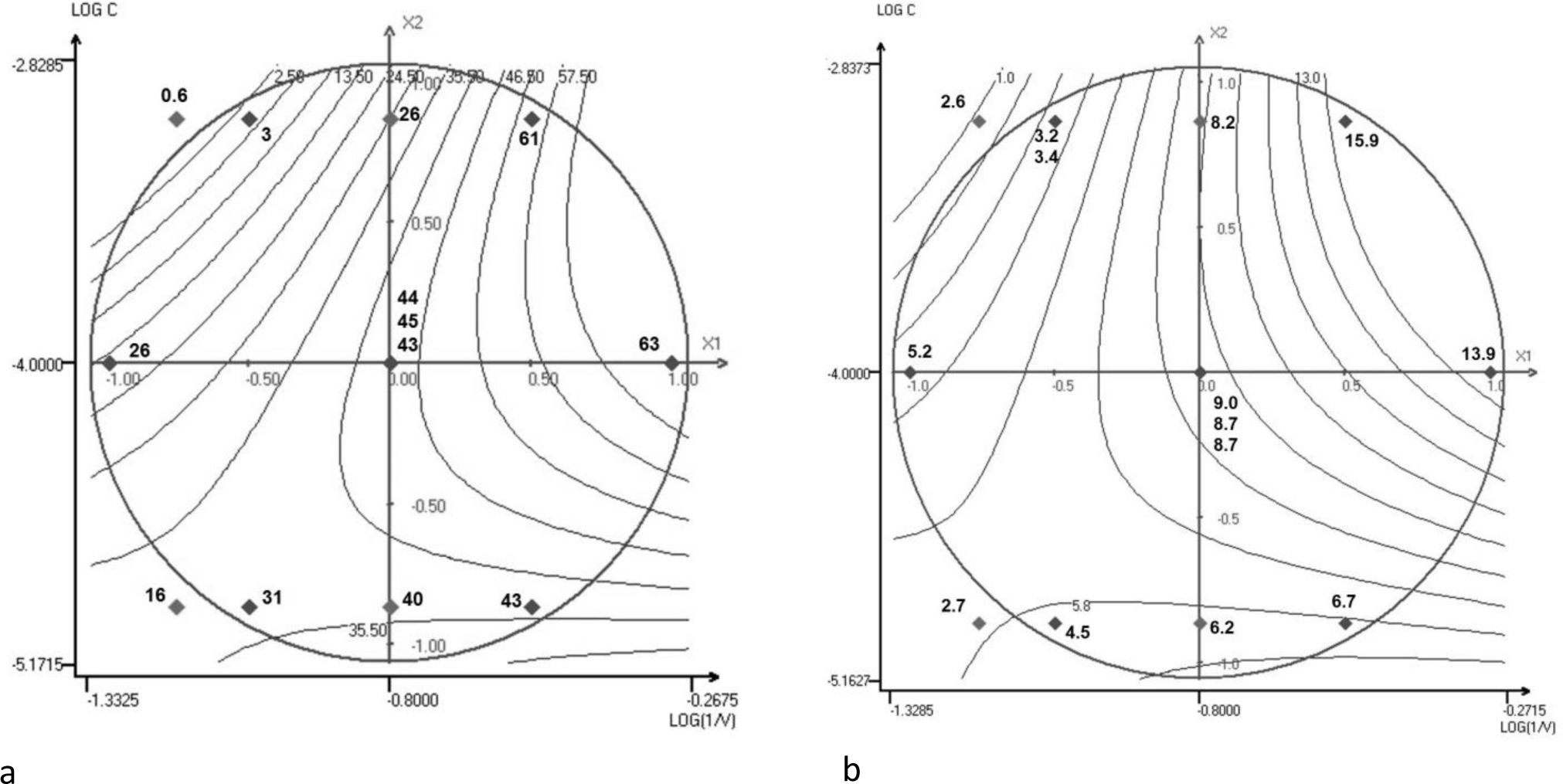
a) Response surface representation for hydroperoxide yield. b) Response surface representation for the extinction coefficient at 233 nm Note: Experimental points are situated in the figure with their experimental values.

The best yields were obtained at intermediate concentrations of methylene blue and with low solution volumes.

It was therefore important to determine whether monomer/dimer/trimer ratios were affected by high methylene blue concentrations. Several authors have suggested that dimer and trimers can be determined from different absorption bands. However, a UV/visible light spectrum analysis for different concentrations of the dye in ethanol (see supplementary information) showed that the ratios of absorption at 654 nm to absorption at 616 nm and 490 nm did not decrease with methylene blue concentration, as reported for methanol (45) but contrary to what is generally observed in water (46). Thus, dimer formation is not favored by high concentrations of methylene blue. In the presence of oxygen and phospholipids, methylene blue gradually loses its color in visible light, resulting in a completely colorless solution. However, at high methylene blue concentrations and large solution volumes, the methylene blue was degraded only very slightly, with absorbance values remaining high from 220 nm to 340 nm (UV) and from 540 to 700 nm. For absorbance at 654 nm (characteristic wavelength of methylene blue illumination for singlet oxygen generation) and different path lengths, 99.9% of the light at this wavelength was found to have been absorbed after a path length of only 1 mm, so almost 90% of the sample received no illumination (see supplementary information).

Similar qualitative behavior was observed for the mass extinction coefficient at 233 nm (Proba.1 < 0.01, Proba.2 =0.059), the two responses being highly correlated (table 6). Indeed, the presence of conjugated dienes is directly related to the amount of hydroperoxide formed (fig. 2b). This amount did not decrease within the limits of the experimental design, implying that conjugated dienes were not significantly degraded through oxidative scission.

For comparison, the unilluminated sample (covered with a sheet of aluminum foil) had a low extinction coefficient (0.279 L.g^−1^.cm^−1^).

#### 3.2.2. Extinction coefficient ratio

The ratio of extinction coefficients at 233 nm/270 nm was described by the model with a better validity than extinction at 233 nm (Proba.1 = 0.033, Proba.2 = 71.9) and its behavior was different (fig. 3). The equation with coded values for log(1/V) and log(C) can be written:

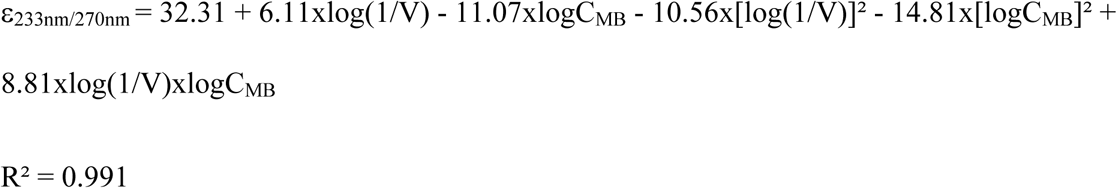

**Figure 3.**
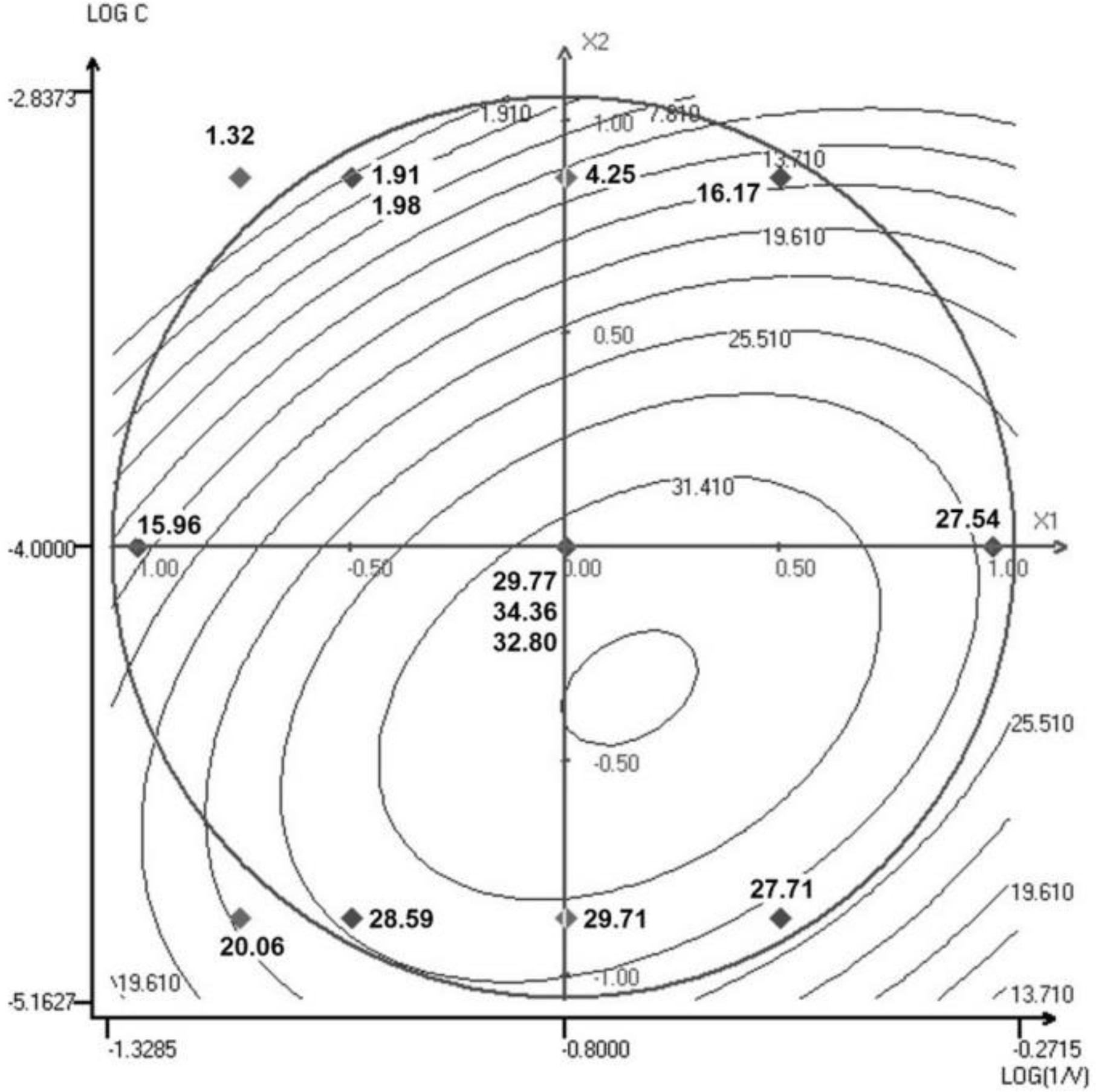
Response surface representation for the ratio of extinction coefficients.

The model could therefore be considered valid for the experimental points used to determine the coefficients, but it was also necessary to check the test points (table 5).

The results for the test points indicated that low extinction ratios displayed greater deviations from the model, and two calculated values (experiments 11 and 13) were significantly different from the experimental values. The model was found to have a high descriptive and moderate predictive value.

The equation confirms that the extinction ratio is inversely correlated with methylene blue concentration (see also table 6). Extinction at 270 nm can be considered characteristic of conjugated trienes, and the change in extinction ratio from poorly to strongly oxidative conditions indicates that these molecules are readily obtained in extreme conditions, with the lowest ratio obtained for a low air volume and high methylene blue concentration. It could be hypothesized that, as only the methylene blue close to the surface is illuminated, these conditions result in a high concentration of singlet oxygen in a small volume. Surface phospholipids are, thus, strongly oxidized, whereas molecules located at deeper positions in the solution are not. Increasing oxygen levels result in greater methylene blue degradation, resulting in the gradual illumination of the entire sample and a homogenization of oxidation mechanisms.

### Size and zeta potential of vesicles

The size of the vesicles formed was determined by drying 1 mL of the phospholipid solution under N_2_, adding 2 mL of deionized water and sonicating the sample for three minutes in pulse mode (5 s ON/10 s OFF). Vesicles were then analyzed by measuring dynamic light scattering (DLS) (table 5).

The Z-average value gives the intensity-weighted harmonic mean size. As this variable was the most stable (see supplementary information), it is selected as the response variable. In terms of mean size, structures smaller than 10 nm across were measured, which may have resulted from the association of a very small number of phospholipids. Based on the size ranges obtained, ultrasonication seems to generate mostly small unilamellar liposomes (47).

The statistical analysis showed that the model obtained for Z-average size was both significant (Proba.1 < 0.01) and valid (Proba.2 =31.9).

The equation for the Z-average size of vesicles was:

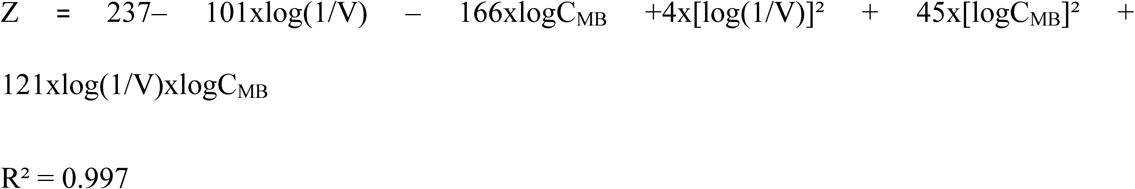

Mean size was not correlated with yield, but was inversely correlated with methylene blue concentration (table 6), as small vesicles were obtained at high methylene blue concentrations, even with only low levels of oxidation. The large decrease in average size with increasing methylene blue concentration and phospholipid solution volume can be deduced from the equation and observed on the response surface (fig. 4).

**Figure 4.**
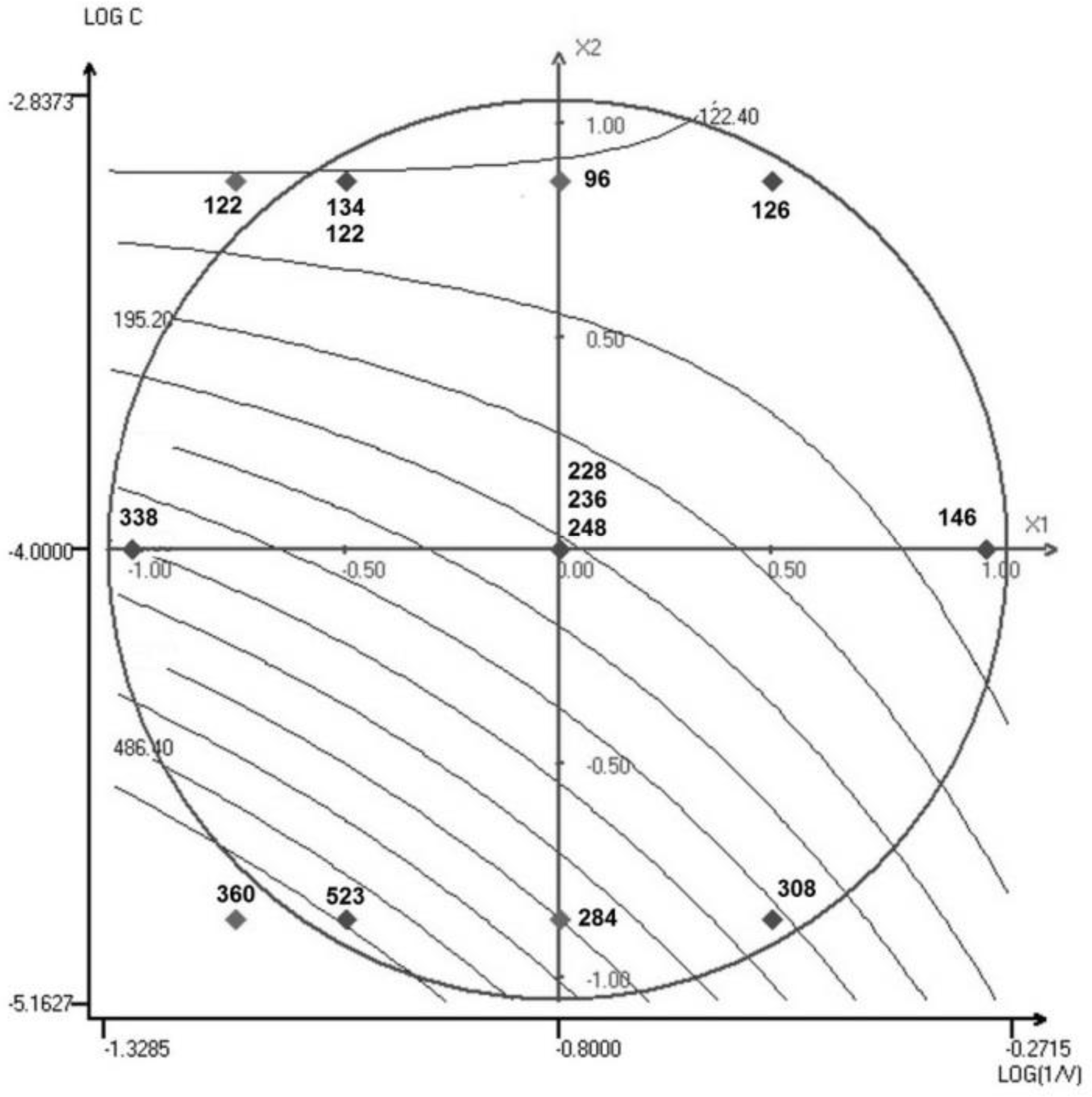
Response surface representation for the Z-average size of vesicles (in nm)

However, due to the instability of some vesicle solutions, the test points gave values either lower than those calculated by the model. The smaller size of the vesicles in oxidized samples may result from mechanical destabilization of the vesicles due to oxidation (48), leading to a partial rupture of vesicles, but it may also indicate that membrane formation requires less energy with oxidized than with native phospholipids. Indeed, increasing the polarity of the fatty acid chains of phospholipids decreases the interaction energy between water and phospholipids. At high concentrations of methylene blue, the volume of solution had a lesser effect. Even if the volume of air is small, a high concentration of methylene blue allows the formation of small vesicles. Despite its presence at a molar concentration well below that of phospholipids (1 mM << ∼25 mM), methylene blue may make a non-negligible contribution to the surface properties of vesicles. As this compound is positively charged, a measurement of the zeta potential of the vesicle solutions is likely to be informative.

Zeta potential can be related to the surface charge of vesicles. The analysis of variance revealed the experimental design to be highly significant (Proba.1 =0.867), whereas the validity of the quadratic model was slightly low (Proba.2 = 3.46).

If the membrane surface consisted solely of the polar heads of phospholipids, then low, mostly negative zeta potentials would be observed, as for vesicles formed without methylene blue and with unoxidized soybean PC only (zeta potential of −8.66 ± 0.42 mV *N*=3). The negativity of the zeta potential in this context is accounted for by the more favorable exposure of the phosphate group than of the choline group (49, 50). When methylene blue is present at a high concentration, without consumption due to a high oxygen level, then zeta potential is positive (fig. 5). This potential can reach values of more than 40 mV and can therefore induce strong electrostatic repulsions between vesicles, limiting aggregation, potentially accounting for the smaller sizes observed. Zeta potential is negative when methylene blue concentration is high but solution volume is low, due to the consumption of methylene blue. Indeed, the degradation of methylene blue under oxidative conditions leads to a loss of its charge (51, 52). This strong contribution of methylene blue to zeta potential, as confirmed by the high correlation coefficient (table 6), results in its positioning at the surface of vesicles, close to the phospholipids, accounting for its preferential reaction with phospholipids rather than with fatty acids. As a cationic molecule, methylene blue adsorbs onto the zwitterionic surface and may change the orientation of the P-N dipole, tilting it away (53). This may increase the expression of the positive charge carried by the choline group. More surprisingly, positive zeta potentials were also observed in the presence of low methylene blue concentrations and small solution volumes. In small volumes, methylene blue is totally degraded and zeta potential depends principally on the conformation of the phosphatidylcholine group. The orientation of the phosphocholine group at the membrane surface may depend on ionic strength(50). Conductivity increases with increasing methylene blue concentration (from about 0.01 mS/cm to 0.1 mS/cm), but the anionic charge on phosphatidyl groups should be more strongly expressed at low concentrations. Oxidation allows more polar fatty acid chains to move toward the external aqueous phase (54, 55), potentially interacting with anionic phosphate groups. Hydroperoxidized fatty acid chains seem to remain in the membrane core (56). Thus, in the absence of photosensitization, oxidation may generate more diverse products interacting preferentially with the phosphate group.

**Figure 5.**
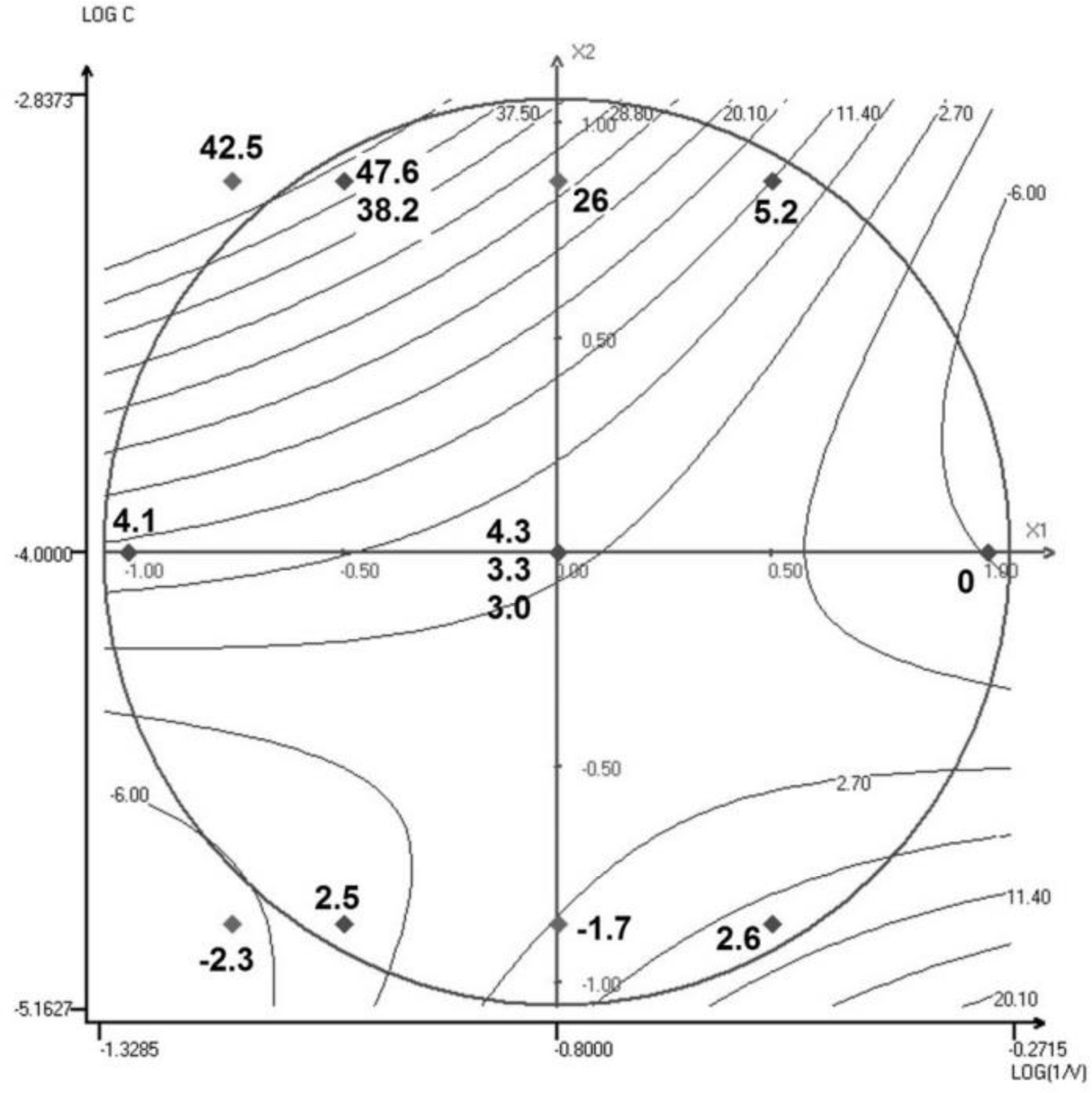
Response surface representation for the zeta potential of vesicles

#### 3.2.3. Permeation studies

KCl is a salt known to permeate liposomes less efficiently than water (10). Changes in the size of liposomes formed from unoxidized phospholipids are therefore more likely to be due to water fluxes than to the movement of KCl. When a solution of vesicles formed with pure dilinoleylphosphatidylcholine (DLiPC) in 0.1 M KCl is dispersed in 2 M KCl, absorbance initially increases considerably, due to light scattering by vesicles. The vesicles are far less permeable to KCl than to water, so water efflux occurs and the vesicle shrinks. An increase in absorbance is observed over recording times of more than 5 s (Abs_0_– Abs_eq_. =-0.072 ± 0.001; Abs_eq_.= 0.585±0.028). A similar absorbance profile is obtained with soybean phosphatidylcholine, but with a lower absorbance and less pronounced variation, suggesting a lower permeability of these vesicles to water (see supplementary information). The main barrier to permeation is generally the highest ordered section of the bilayer. Unsaturation leads to lower chain-ordering values and an increase in membrane fluidity, potentially leading to transient defects or pores, increasing water permeation. Soybean phosphatidylcholine cans fewer unsaturated bonds and is therefore, less permeable to water (5–7).

The absorbance of many samples did not change significantly over the time course of the permeation test, making it impossible to model the ΔA parameter correctly. However, certain types of behavior emerged. The sample corresponding to experiment 14 (tables 4 and 5) oxidized with the lowest methylene blue concentration and a small volume of air, with a low extinction coefficient at 233 nm and a yield (2.72 and 16.25%, respectively) similar to that for soybean PC. Vesicles formed with phospholipids oxidized at the lowest methylene blue concentration and a medium–high air volume mostly displayed a decrease in absorbance over time (samples 4, 5 and 12), indicating swelling and an increase in KCl permeation. Samples with medium-high methylene blue concentrations but low air volumes (2 and 13) formed small vesicles with a high absorbance that increased over time, indicating shrinkage due to water efflux. This high absorbance may also be partly due to the presence of unconsumed methylene blue rather than purely the result of scattering. Phospholipids oxidized with medium-high air volumes and medium-high methylene blue concentrations (samples 1, 3, 7, 8, 9, 11), which have had extinction coefficients at 233 nm (>8 L.g^−1^.cm^−1^), formed vesicle suspensions with a low absorbance that did not vary over time, indicative of a total absence of swelling/shrinkage behavior. This low absorbance can be linked to the small size of the vesicles and the absence of variation may be due to the high level of membrane permeability, as also observed for samples 6 and 10. This behavior, despite low hydroperoxide yields and low 233 nm extinction ratios, may be due to the high concentration of methylene blue, suggesting that this molecule can promote membrane permeability in the presence of low-level oxidation. The surface response corresponding to the final absorbance A_inf_ (fig. 6) can be obtained, to assess the extent to which it is linked to initial vesicle size or methylene blue concentration. Statistical analysis showed the model to be highly significant (Proba.1 <0.01), with good validity (Proba.2 =84.1).

**Figure 6.**
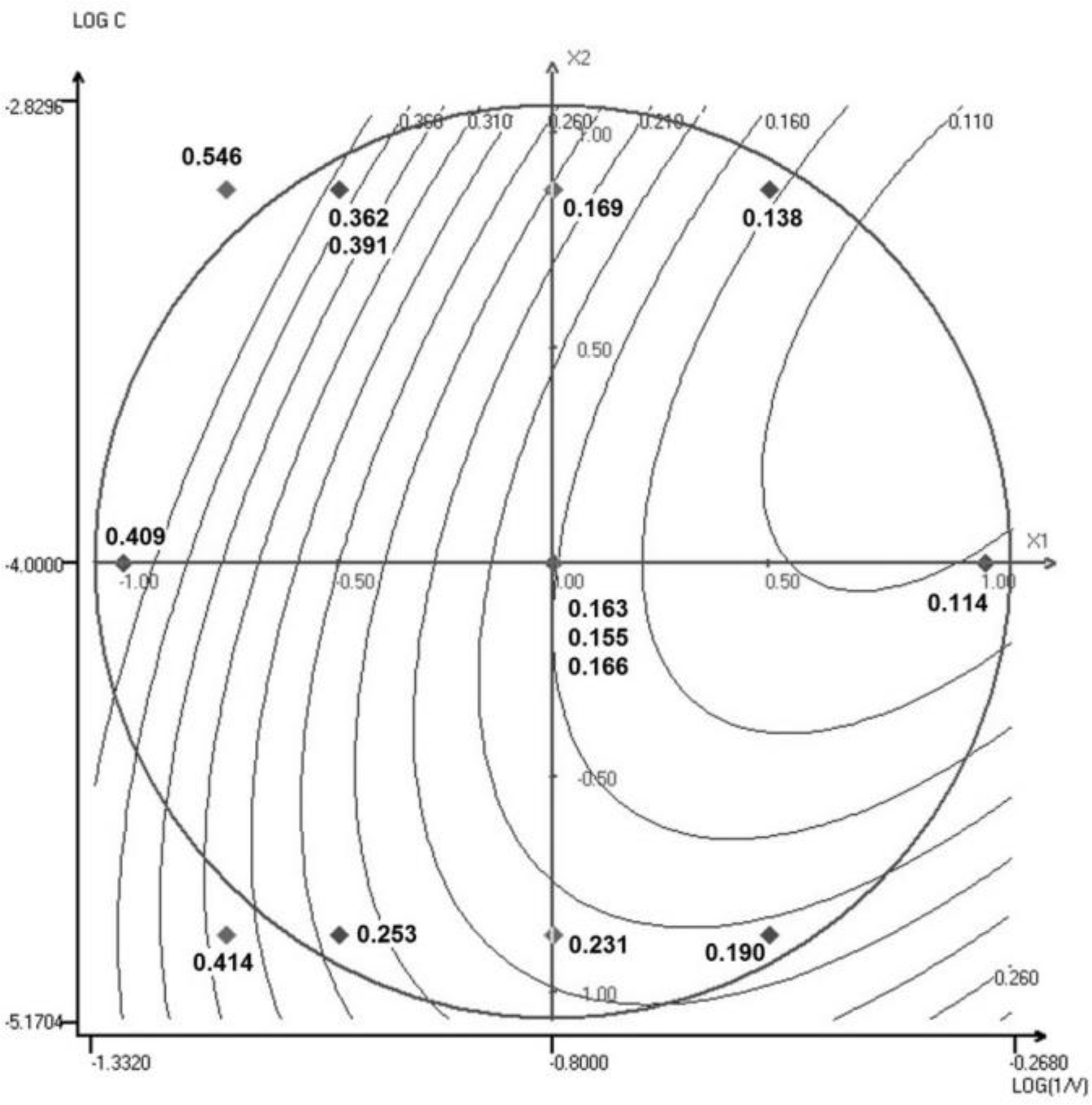
Response surface representation for the final absorbance of vesicles

The equation determined for final absorbance was:

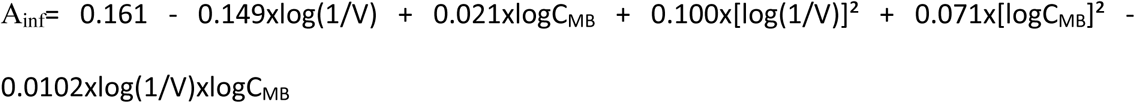

The test points displayed only limited divergence from the calculated values, so the model describes experimental values well, but also has predictive power. This predictive power could be improved by increasing the degrees of freedom. The shape of the response surface was different from that for vesicle size (fig. 6). Even if the positive contribution of methylene blue to absorbance must be taken into account, this figure, the equation above and table 6 indicate that final absorbance is correlated principally with solution volume, whereas vesicle size is essentially inversely correlated with methylene blue concentration. Both yield and extinction coefficients are inversely correlated with final absorbance, and are therefore not entirely dependent on initial size but on the hydroperoxide content of the phospholipids forming the vesicle membrane, determining its ability to swell/shrink.

## 4. Conclusion

Methylene blue favored interaction with soybean phospholipids. Used as a tool for monitoring the photosensitized oxidation of phospholipids, methylene blue could improve our understanding of membrane modifications at low to medium levels of oxidation. Its influence on the surface charge and size of phospholipid vesicles can be controlled through response surface methods and could be used to adapt the stability and transport properties of liposomes.

## Author Contribution section

J-F. F. designed and performed research, analyzed data and wrote the manuscript.

M. C. contributed analytical tools, analyzed data.

A. C. contributed analytical tools, analyzed data.

Z. M. designed research, analyzed data.

### Acknowledgments

The authors thank the Occitanie region for financial support through the SMON-FERT RECH project, grant no. 1405981.

The authors thank Mr. Marc Vedrenne of the University of Toulouse for assistance with NMR analysis and Dr Aurélie BEAL and Prof. Roger PHAN TAN LUU from NEMRODW for scientific assistance.

## Supporting information

One file containing statistical processing tables of the experimental design, details of NMR analysis, absorbance variations of methylene blue solutions with concentration and path length, details of dynamic light scattering measurements and permeation studies.

